# CodY controls the SaeR/S two-component system by modulating branched-chain fatty acid synthesis in *Staphylococcus aureus*

**DOI:** 10.1101/2024.05.03.592463

**Authors:** Shahad Alqahtani, Dennis A. DiMaggio, Shaun R Brinsmade

## Abstract

*Staphylococcus aureus* is a Gram-positive, opportunistic human pathogen that is a leading cause of skin and soft tissue infections and invasive disease worldwide. Virulence in this bacterium is tightly controlled by a network of regulatory factors. One such factor is the global regulatory protein CodY. CodY links branched-chain amino acid sufficiency to the production of surface-associated and secreted factors that facilitate immune evasion and subversion. Our previous work revealed that CodY regulates virulence factor gene expression indirectly in part by controlling the activity of the SaeRS two-component system. While this is correlated with an increase in membrane anteiso-15:0 and −17:0 branched-chain fatty acids (BCFAs) derived from isoleucine, the true mechanism of control has remained elusive. Herein, we report that CodY-dependent regulation of SaeS sensor kinase activity requires BCFA synthesis. During periods of nutrient sufficiency, BCFA synthesis and Sae TCS activity is heavily suppressed by CodY-dependent repression of the *ilv-leu* operon and the isoleucine-specific permease gene *brnQ2.* In a *codY* null mutant, which simulates extreme nutrient limitation, de-repression of *ilv-leu* and *brnQ2* directs the synthesis of enzymes in redundant de novo and import pathways to catalyze the production of BCFA precursors. Overexpression of *brnQ2* independent of CodY is sufficient to increase membrane anteiso BCFAs, Sae-dependent promoter activity, and SaeR∼P levels. Our results further clarify the molecular mechanism by which CodY controls virulence in *S. aureus*.

**IMPORTANCE:** Expression of bacterial virulence genes often correlates with the exhaustion of nutrients, but how the signaling of nutrient availability and the resulting physiological responses are coordinated is unclear. In *S. aureus,* CodY controls the activity of two major regulators of virulence – the Agr and Sae two-component systems – by unknown mechanisms. This work identifies a mechanism by which CodY controls the activity of the sensor kinase SaeS by modulating the flux of anteiso branched-chain amino acids to the membrane. Understanding the mechanism adds to our understanding of how bacterial physiology and metabolism are linked to virulence and underscores the homeostatic nature of virulence. Understanding the mechanism also opens potential avenues for targeted therapeutic strategies against *S. aureus* infections.

## Introduction

*Staphylococcus aureus* is a Gram-positive bacterium found commonly on human skin and in the anterior nares. Up to 30% of the human population may be stably colonized by this pathogen without experiencing any symptoms. As an opportunistic human pathogen, *S. aureus* is the leading cause of skin and soft tissue infections. These infections can progress to devastating invasive infections including infectious endocarditis, osteomyelitis, and sepsis (1, 2). Once confined to healthcare settings, nosocomial infections of patients occurred in hospitals for decades. Resistance to methicillin (MRSA) and other antimicrobials compounded the problem. More troubling is the relatively recent spread into communities (CA-MRSA). Over the past decade, CA-MRSA has killed tens of thousands of people (3), and CA-MRSA infections cost the U.S. healthcare system between $560 million and $2.7 billion in annual treatment-associated costs (4).

*S. aureus* can infect nearly every organ of the body. The success of *S. aureus* as a pathogen has been attributed to its vast repertoire of surface-associated and secreted virulence factors that enhance host colonization and facilitate evasion from the host immune response (5). However, from a resource and energy conservation prospective, the production of all virulence factors simultaneously is likely to be costly to the bacterium. Therefore, the expression of the virulence factors is tightly regulated by a network of transcription factors, two-component systems, and small regulatory RNAs (6–8). As a result, the bacterium licenses the production of only those factors relevant to a given infection niche. Interfering with the regulation and production of virulence factors may be our best hope for combating pathogens resistant to available antimicrobial therapies (9).

The production of virulence factors in *S. aureus* is controlled in large part by the Sae two-component system (TCS). Sae is known to regulate the expression of over 20 surface-associated and secreted virulence genes such as *hla* (ɑ-hemolysin), *coa* (coagulase), *nuc* (thermonuclease), and proteases (10–13). The *sae* locus consists of a weak constitutive P3 promoter that drives the expression of the response regulator gene *saeR* and the histidine kinase gene *saeS.* In addition, an inducible P1 promoter controls the transcription of two auxiliary genes *saeP* and *saeQ* (14–16) along with *saeRS*. SaeS is an atypical histidine kinase. Unlike typical histidine kinases that contain a large extracellular domain that binds a ligand to transduce signals across the membrane, SaeS belongs to the intramembrane family of histidine kinases (17) that lack this domain. Indeed, SaeS only has a nine amino acid residue extracellular-facing peptide that links two transmembrane helices. Together, this N-terminal domain constrains SaeS kinase activity in the cytosol at the C-terminus (18). In the presence of neutrophil-produced factors such as human neutrophil peptides 1, 2, and 3 (HNP1-3), SaeS is phosphorylated on a conserved histidine residue by an unknown mechanism. The phosphoryl group is then transferred to a specific, conserved aspartate residue in SaeR (to generate SaeR∼P). SaeP is a lipoprotein, and SaeQ is predicted to be a transmembrane protein. Previous work has shown that SaeP and SaeQ form a complex with SaeS and act to stimulate the phosphatase activity of SaeS toward SaeR by an unknown mechanism. This lowers cellular SaeR∼P levels and returns the Sae TCS to the pre-induced state (19). SaeR∼P binds directly to dozens of targets with variable affinity (10, 20). This includes the *sae* P1 promoter, triggering the production of SaeP and SaeQ, creating a negative feedback loop to prevent Sae activity from becoming unlimited.

Branched-chain fatty acids (BCFAs) are the most abundant membrane fatty acids in staphylococcal membranes and we previously reported that BCFA levels are correlated with SaeS kinase activity (21). The synthesis of BCFAs in *S. aureus* begins with the amino acids isoleucine, leucine, and valine (ILV). These amino acids can be imported into the cell through specific transporters BrnQ1, BrnQ2, and BcaP, or synthesized within the cell under certain conditions (22–25). After ILV import and transamination by the enzyme IlvE, the resulting branched-chain α-keto acids undergo oxidative decarboxylation to form branched-chain carboxylic acids (BCCAs) that are subsequently activated to their acyl-CoA derivatives. Under laboratory conditions, this series of reactions is catalyzed by the branched-chain α-keto acid dehydrogenase complex (BKDH) and associated coenzymes (26). The acyl-CoAs are then primed by FabH enzyme, which catalyzes their condensation with malonyl-ACP. The resulting β-ketoacyl-ACP is further elongated by the type II fatty acid synthase (FASII) before being incorporated into phospholipids (27). In *S. aureus*, the most abundant BCFAs in the cell membrane are iso (*i*) and anteiso (*a*) fatty acids derived from isoleucine, specifically *a*15:0 and *a*17:0 (22, 28).

CodY (AKA, **C**ontroller **o**f ***d****pp*) was first discovered in *Bacillus subtilis* as a repressor of dipeptides permease (*dpp*), and then later a variety of permeases, biosynthetic genes, and alternative nutrient processing enzyme genes during rapid growth (i.e., metabolic genes) (29–31). We now know that CodY is a global regulatory protein that controls the expression of dozens of genes in many low G+C Gram-positive bacterial genera, including *Bacillus, Listeria, Stapylococcus, Clostridioides, Lactococcus,* and *Streptococcus* (32–42). In pathogens like *S. aureus*, CodY also regulates the expression of virulence genes (39, 40, 43). As such, CodY serves as an important linkage between metabolism and pathogenic potential (44). CodY is activated as a DNA-binding protein when bound to ILV and GTP (39, 40, 45, 46). CodY interacts with a site-specific DNA sequence essential for binding target genes (AATTTTCWGAAAATT) originally determined in *Lactococcus lactis* and validated in *S. aureus* (33, 39). CodY prioritizes gene expression based in part on DNA-binding activity controlled by availability of ILV and affinity of CodY for its many sites to facilitate the bacterium’s adaptation to its environment (34, 47–50). For many of the metabolic genes, CodY directly represses gene expression at target promoters. In contrast, CodY control of virulence in *S. aureus* is indirect via two major regulators of virulence – the Agr quorum sensing system and the Sae TCS (21, 39, 51, 52). This control mediates the production of secreted digestive enzymes and cytotoxins to replenish nutrients during infection when amino acid and energy reserves are low. In support, knocking out CodY results in hypervirulence in mouse models of skin and soft tissue infection (SSTI) and necrotizing pneumonia (53). At least for SSTI, this is due to CodY-dependent regulation of *hla* (*α*-toxin) (21). Although CodY adjusts the expression of the *sae* locus by direct and indirect mechanisms at the *sae* P1 promoter (52) we previously showed that this transcriptional control is dispensable. Rather, CodY controls the activity of the SaeS kinase by an unknown mechanism correlated with changes to membrane fatty acid content (21). Herein, we report that CodY-deficient strains disrupted for de novo ILV synthesis and isoleucine import exhibit reduced Sae-dependent gene expression and SaeS kinase activity. This is correlated with lower levels of anteiso 15:0 and 17:0 branched-chain fatty acids derived from isoleucine. This low Sae TCS activity phenotype is complemented genetically, and also chemically by supplementing mutant cells with exogenous branched-chain fatty acids. We show that overexpression of the isoleucine-specific permease *brnQ2* is sufficient to activate Sae TCS activity in WT cells during laboratory growth, and this increased activity is correlated with the level of the anteiso 15:0 and 17:0 BCFAs in the membrane. Our results further clarify the mechanism by which CodY controls a major virulence regulator to optimize virulence factor production during nutrient depletion.

## Results

### Branched-chain fatty acid synthesis is required to upregulate Sae TCS activity when CodY activity is reduced

We previously demonstrated that BCFAs are essential for Sae TCS activity, and BCFA content is increased in Δ*codY* mutant cells (21). We wondered whether *a*15:0 BCFAs are required to upregulate Sae TCS activity when *codY* is knocked out. To test this, we took advantage of a mutant strain that synthesizes *i*14:0 BCFAs for growth but not *a*15:0 BCFAs to promote Sae activity and performed an epistasis experiment using the Sae-dependent *saeP1-gfp* reporter fusion to indirectly measure Sae activity. The strain (11*lpdA mbcS1*) is devoid of BKDH activity and overexpresses a methylbutryl-CoA synthetase MbcS that activates exogenously supplied branched carboxylic acids (i.e., isobutyric acid; *i*C_4_) (54, 55). As expected, when we knocked out *codY, saeP1-gfp* promoter activity increased 2-fold during growth in rich, complex medium (i.e., tryptic soy broth [TSB]). We measured very low promoter activity in the 11*lpdA mbcS1* mutant (**Fig 1**, compare *lpdA mbcS1* and *codY* to WT). When we knocked out *codY* in the *lpdA mbcS1* background, promoter activity was indistinguishable from that measured in the *lpdA mbcS1* strain. Promoter activity was restored when we supplemented 11*lpdA mbcS1* double mutant and Δ*lpdA mbcS1* 11*codY* triple mutant cells with 0.5 mM of the individual short, branched-chain carboxylic acids *a*C_5_ (2-methylbutyric acid) and *i*C_4_ (isobutyric acid) but not *i*C_5_ (3-methylbutyric acid). We saw no additive or negative effect when we provided a mixture of all three carboxylic acid precursors. These data indicate that 11*lpdA mbcS1* is epistatic to Δ*codY,* and BCFAs are required for CodY-dependent up-regulation of Sae TCS activity. The effect is specific for some but not all BCFAs, but we did not test whether there was a preference for anteiso BCFAs over iso BCFAs.

**Figure 1.**
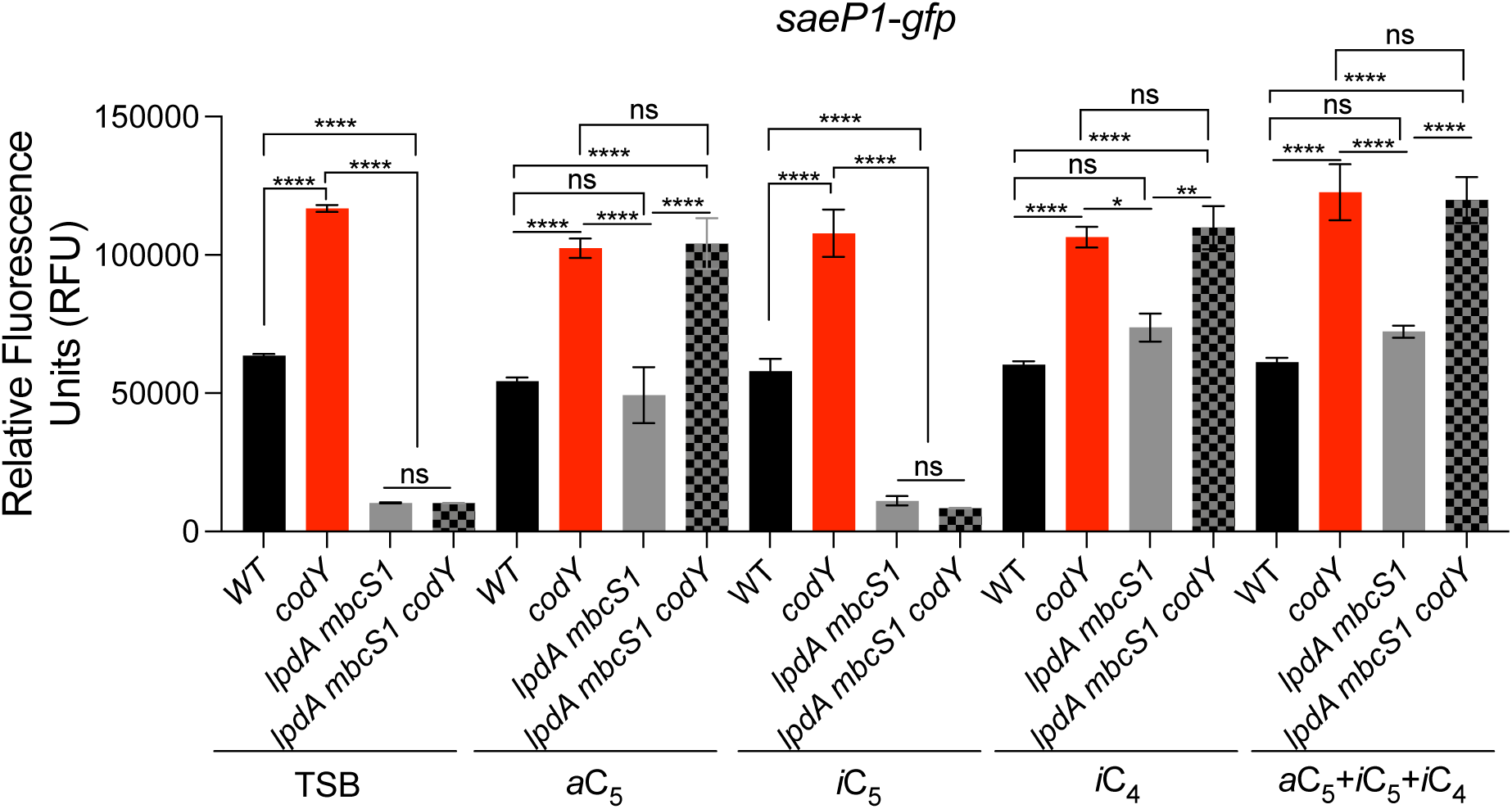
CodY control of Sae TCS activity requires BCFAs. The indicated USA300 LAC strains harboring the *saeP1-gfp* reporter fusion were grown in TSB or TSB supplemented with 0.5 mM BCFA precursors *a*C_5_ (2-methylbutyric acid), *i*C_5_ (3-Methylbutyric acid), and *i*C_4_ (isobutyric acid) for 16 h overnight. Data are plotted as mean ± SEM of three biological replicates. \**p* < 0.05, \*\**p* < 0.01, \*\*\**p* < 0.001, \*\*\*\**p* < 0.0001, one-way ANOVA with Tukey’s multiple comparison test within each treatment group. Ns, not significant.

### *De novo* amino acid biosynthetic or salvaging pathways are required for CodY-dependent regulation of Sae TCS activity

During laboratory growth in ILV-replete medium, BCFAs are synthesized from ILV, and the membrane fatty acid composition depends in part on the levels of exogenous amino acids and selectivity of the BKDH complex for isoleucine (22). CodY represses genes coding for the enzymes that direct *de novo* ILV biosynthesis (*ilv-leu*), and the ILV permeases *brnQ1* and *brnQ2.* BrnQ1 transports all three branched-chain amino acids, whereas BrnQ2 transports isoleucine specifically (23, 39, 48). We hypothesized that CodY restricts BCFA synthesis during conditions of BCAA sufficiency and keeps Sae TCS activity low. During BCAA limitation, reduced CodY activity would result in upregulation of ILV import and synthesis pathways, leading to increased branched-chain ɑ-keto acids levels that would feed branched-chain fatty acid synthesis and upregulate Sae activity when incorporated into the membrane. To begin to test this hypothesis, we disrupted *de novo* ILV biosynthesis or importer genes in the 11*codY* mutant and monitored Sae activity indirectly using the Sae-dependent *nuc-gfp* reporter fusion (48, 56) during growth in chemically defined medium (CDM). We used CDM to simplify the experimental conditions, as TSB contains peptides as well as free amino acids. As expected, *nuc-gfp* promoter activity increased by ∼4-fold in the Δ*codY* mutant. Deleting *ilvD,* or disrupting *brnQ1* or *brnQ2* in the Δ*codY* mutant background did not significantly reduce *nuc-gfp* activity (**Fig 2A**, compare double mutants to the *codY* mutant). However, synthesis and import are redundant pathways for maintaining ILV levels and BCFA precursor pools. Therefore, we made triple mutants. While disrupting *ilvD* and *brnQ1* in the Δ*codY* mutant did not alter *nuc-gfp* promoter activity, when we knocked out both *ilvD* and *brnQ2* in the Δ*codY* mutant we measured a 2-fold drop in promoter activity (**Figure 2A**). We did not knock out additional permease genes in this background (i.e., *bcaP*) as this would likely result in a confounding growth defect in CDM. We then analyzed the membrane fatty acid content of the 11*codY* 11*ilvD* and 11*codY* 11*ilvD* 11*brnQ2* strains using gas chromatography analysis of fatty acid methyl esters (GC-FAME). Sae-dependent promoter activity was correlated with anteiso BCFA levels. Specifically, we measured a decrease in *a*15:0 and *a*17:0 BCFAs in the triple mutant. *nuc-gfp* promoter activity, BCFA content, and secreted nuclease production phenotypes were complemented chemically when we included a mixture of the carboxylic acid precursors in the medium or genetically when we expressed *brnQ2^+^*ectopically on the multicopy plasmid pRMC2 (**Figure 2B-D**).

**Figure 2.**
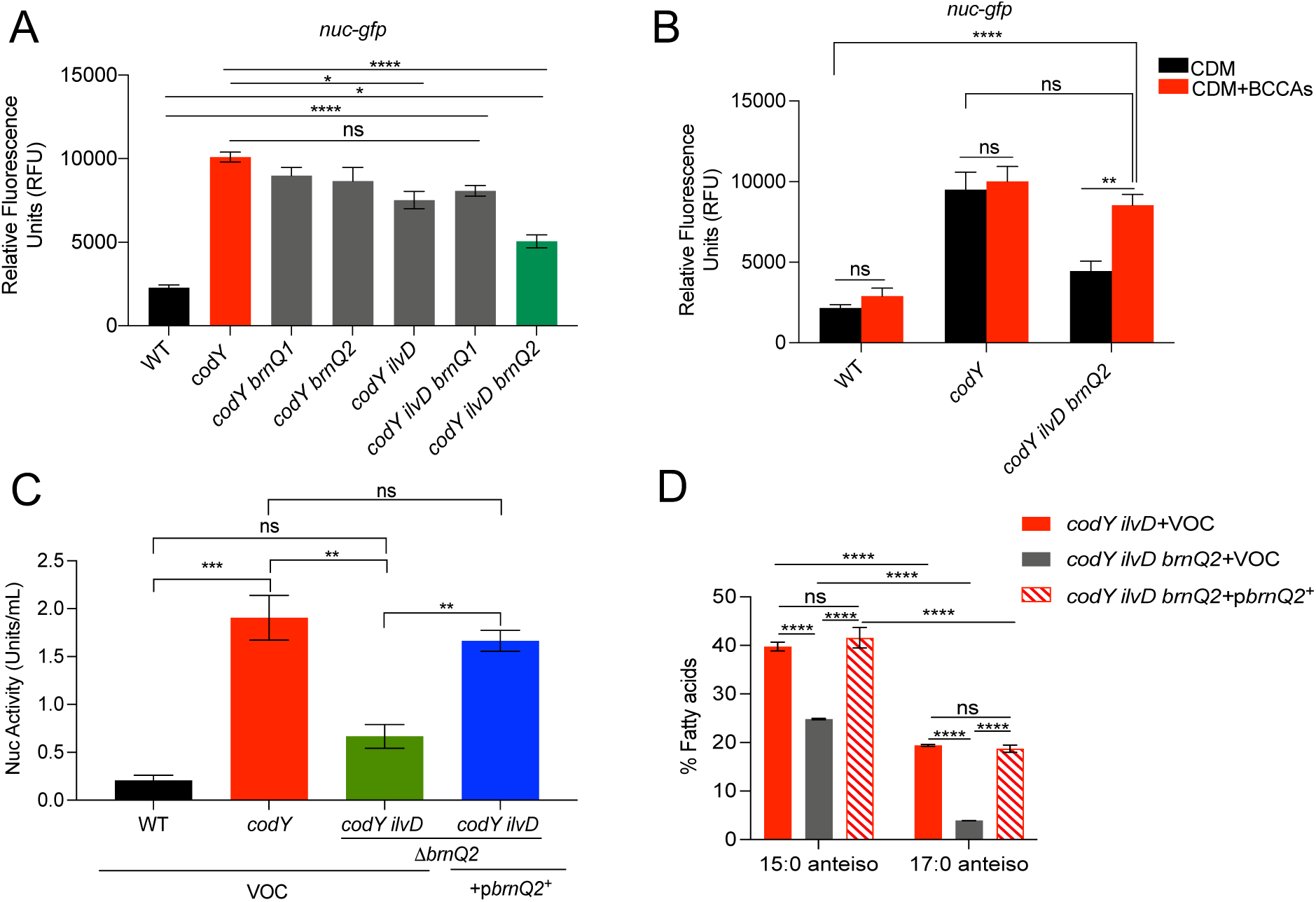
CodY control of Sae TCS activity requires *de novo* α-keto acid synthesis or isoleucine import. (A and B) The indicated LAC strains were grown for 16 h in chemically defined medium (CDM), at which time *nuc-gfp* promoter activity was measured. (C and D) The indicated strains were grown to exponential phase in CDM, at which time secreted nuclease activity was measured and membrane fatty acid content was analyzed using GC-FAME. For all panels, data are plotted as mean +/- SEM of at least three independent experiments. \**p* < 0.05; \*\**p* < 0.01; \*\*\**p* < 0.001, \*\*\*\**p* < 0.0001. Significance was determined using one-way ANOVA with repeated measures and Tukey posttest for (A and C) or Two-way ANOVA for (B and D). BCCAs, 0.5 mM each of *a*C_5_, *i*C_4_, *i*C_5_; ns, not significant.

If CodY controls the Sae TCS by controlling the synthesis of BCFAs, then uncoupling *brnQ2* or *ilv-leu* expression from CodY regulation and overexpressing the gene(s) in a WT background should phenocopy the 11*codY* mutant for heightened Sae activity. To test this, we expressed *brnQ2^+^* from the inducible P*_tet_* promoter in WT cells and monitored Sae TCS activity using the *saeP1-gfp* reporter fusion. As expected, promoter activity increased > 2-fold in the 11*codY* mutant relative to WT; the fusion is Sae-dependent (**Fig S1**). Compared to the vector control, we observed a dose-dependent increase of *saeP1-gfp* promoter activity as we increased the concentration of anhydrotetracycline in the medium. At higher levels of aTc (e.g., 25 ng ml^-1^), promoter activity exceeded that measured in the *codY* mutant (**Fig 3A** and **Fig S1**). This increase in SaeS activity was not observed in WT cells containing *saeP1-gfp* and the pRMC2 vector-only control (**Fig 3A)**. Again, increased promoter activity was correlated with a significant increase in 15:0 anteiso BCFAs. Although not significant, we saw an increase in 17:0 anteiso BCFA levels as well (**Fig 3B)**. These data strongly suggest that de novo amino acid biosynthetic or salvaging pathways are required to upregulate Sae activity when CodY activity is reduced. Further, overexpressing *brnQ2* results in the same activation phenotype, implying that BCFA flux to the membrane, but not differential production of membrane proteins under CodY control is required. The mechanism is consistent with our previous work, revealing that Sae TCS activity is affected by isoleucine-derived phospholipids (21).

**Figure 3.**
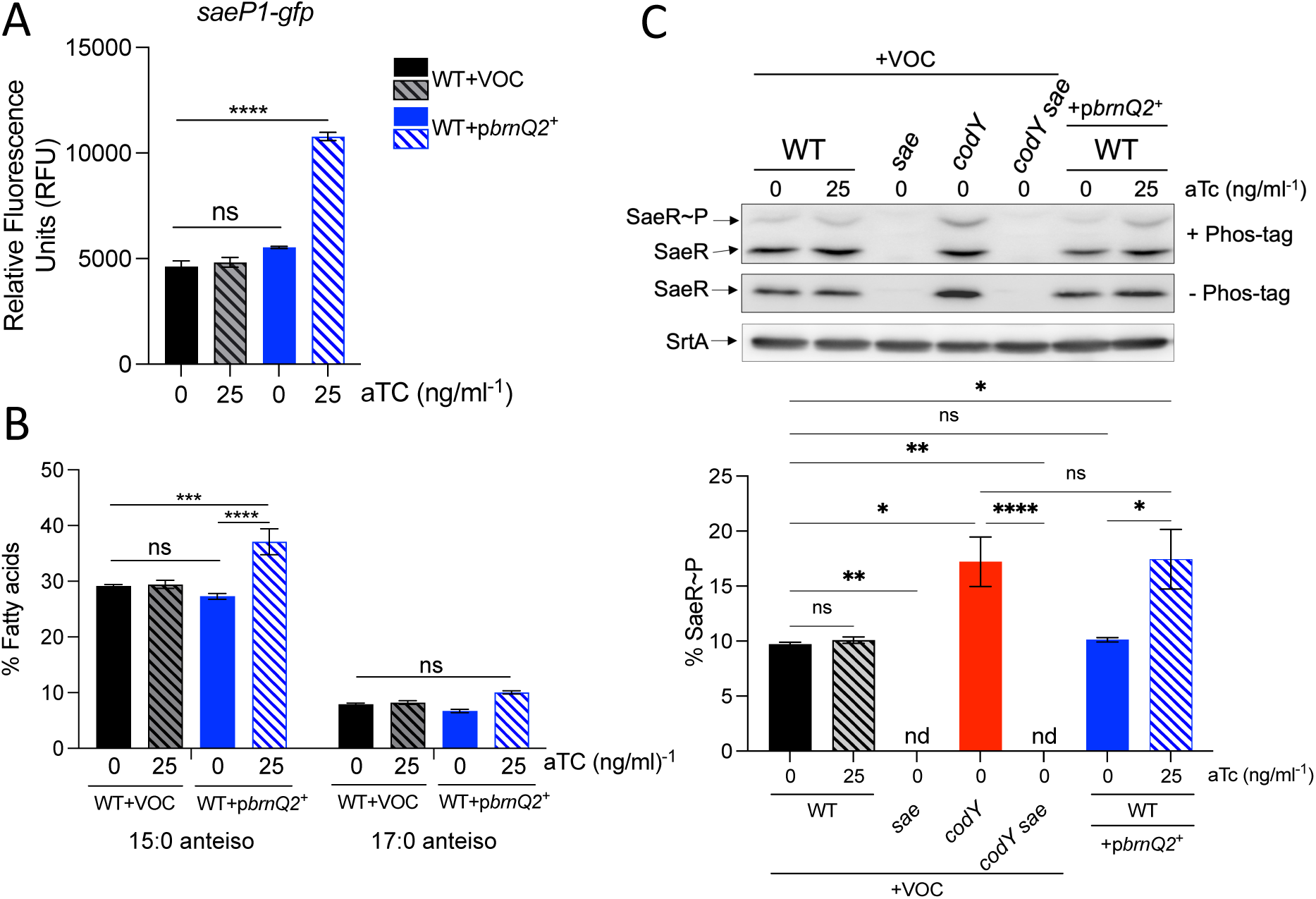
Overexpressing the isoleucine permease gene *brnQ2* results in higher levels of anteiso BCFAs and SaeS activity. Strains were grown to exponential phase in CDM medium with and without anhydrotetracycline (aTC). Cells were collected and (A) *saeP1-gfp* promoter activity (B) membrane fatty acid composition and (C) intracellular SaeR protein species were measured. SaeR was detected in SDS-polyacrylamide gels without and with Phos-Tag reagent using polyclonal antibodies raised against SaeR. SrtA was detected as a loading control (upper; representative gel). Densitometric quantification was performed using ImageJ (lower). Data are plotted as mean +/- SEM of at least three independent experiments. \**p*<0.05, \*\**p*<0.01, \*\*\**p*<0.001, \*\*\*\**p*<0.0001, one-way ANOVA with Tukey’s multiple comparison test. ns, not significant; nd, not detected.

Unphosphorylated SaeR does not bind target promoters. Rather, activated SaeR (SaeR∼P) binds to stimulate gene transcription (20). Further, each histidine kinase-response regulator pair functions autonomously. Crosstalk between noncognate HK-RR pairs rarely occurs. Therefore, SaeS kinase activity solely determines the cellular levels of SaeR∼P (57). To test directly whether overexpressing *brnQ2* increases SaeS kinase activity, we used Phos-Tag electrophoresis (58) and Western blotting with SaeR antisera to measure the relative levels of SaeR and SaeR∼P in WT cells expressing *brnQ2* from the inducible P*_tet_* promoter. As expected, the fraction of SaeR∼P was ∼10% in WT cells in the pre-induced state; this increased to ∼20% in the *codY* mutant. Compared to vector controls, when we overexpressed *brnQ2^+^* in WT cells, SaeR∼P levels were essentially identical to those measured in the *codY* mutant. This was correlated with a modest increase in total SaeR levels, likely due to autoregulation of *sae* expression via the *sae* P1 promoter (**Fig 3C** and **Fig S2**) (21). Thus, SaeS kinase activity increases when isoleucine import via BrnQ2 is increased.

## Discussion

SaeS is a member of the intramembrane family of histidine kinases for which relatively little is known. A defining feature of these kinases is the lack of a large extracellular ligand/signal-binding domain that perceives environmental stimuli (17). Rather, a short peptide links two transmembrane helices. The prevailing view for SaeS is that the conformation of the N-terminal domain controls the activity of the C-terminal cytoplasmic kinase domain (18). In this context, it is conceivable that any signal, internal or external, can influence SaeS kinase activity to upregulate the expression of virulence genes by altering the membrane environment. Indeed, human neutrophil peptides, which are produced by immune cells and perforate the bacterial membrane, induce Sae TCS activity by an unknown mechanism. We previously discovered that CodY controls nearly all *S. aureus* virulence genes indirectly in part by controlling the activity of the Sae TCS (48, 52). However, the mechanism by which CodY upregulates Sae activity has remained unclear. Herein, we show that the upregulation of Sae TCS activity in *codY* mutant cells depends on anteiso BCFA synthesis using exogenous or endogenous branched-chain carboxylic acid precursors. Endogenous synthesis of the precursors is controlled by both de novo synthesis via the ILV biosynthetic pathway or by ILV import and transamination by IlvE. The result is an increase in SaeS kinase activity and cellular SaeR∼P levels to stimulate Sae-dependent promoters.

Our findings build upon previous work by Pendleton *et al.,* demonstrating the essential role of BCFAs in Sae TCS activity and informing our working model (**Fig 4**). The question that remains is how exactly BCFAs affect the SaeS signaling complex. As described above, the prevailing view is that the conformation of the transmembrane portion of SaeS controls activation, but how HNPs alter the conformation is an open question. A growing body of research suggests that the transmembrane helices connect the IM-HKs to additional membrane proteins. SaeS was reported to be localized in functional membrane microdomains (FMMs), regions of low fluidity in an otherwise fluid membrane (59). One possibility is that anteiso BCFAs specifically facilitate the diffusion of an activator protein through the bilayer into the FMM, or facilitate SaeS diffusion out of the FMM to interact with the activator protein. On the other hand, anteiso BCFAs may facilitate the dispersal of inhibitor proteins out of the SaeS signaling complex. Indeed, SaeS is known to interact with auxiliary membrane proteins SaeP and SaeQ, which induce SaeS phosphatase activity by an unknown mechanism. This returns Sae TCS activity to its pre-induced state. However, at least for CodY-dependent regulation, SaeP and SaeQ are not involved, as CodY control is retained in a SaePQ-deficient strain (21). We cannot exclude the possibility that HNP-mediated induction occurs in a SaePQ-dependent manner. Another variation on this theme is that other protein-protein interactions are affected in an anteiso BCFA-dependent manner that somehow indirectly affects the activity of SaeS.

**Figure 4.**
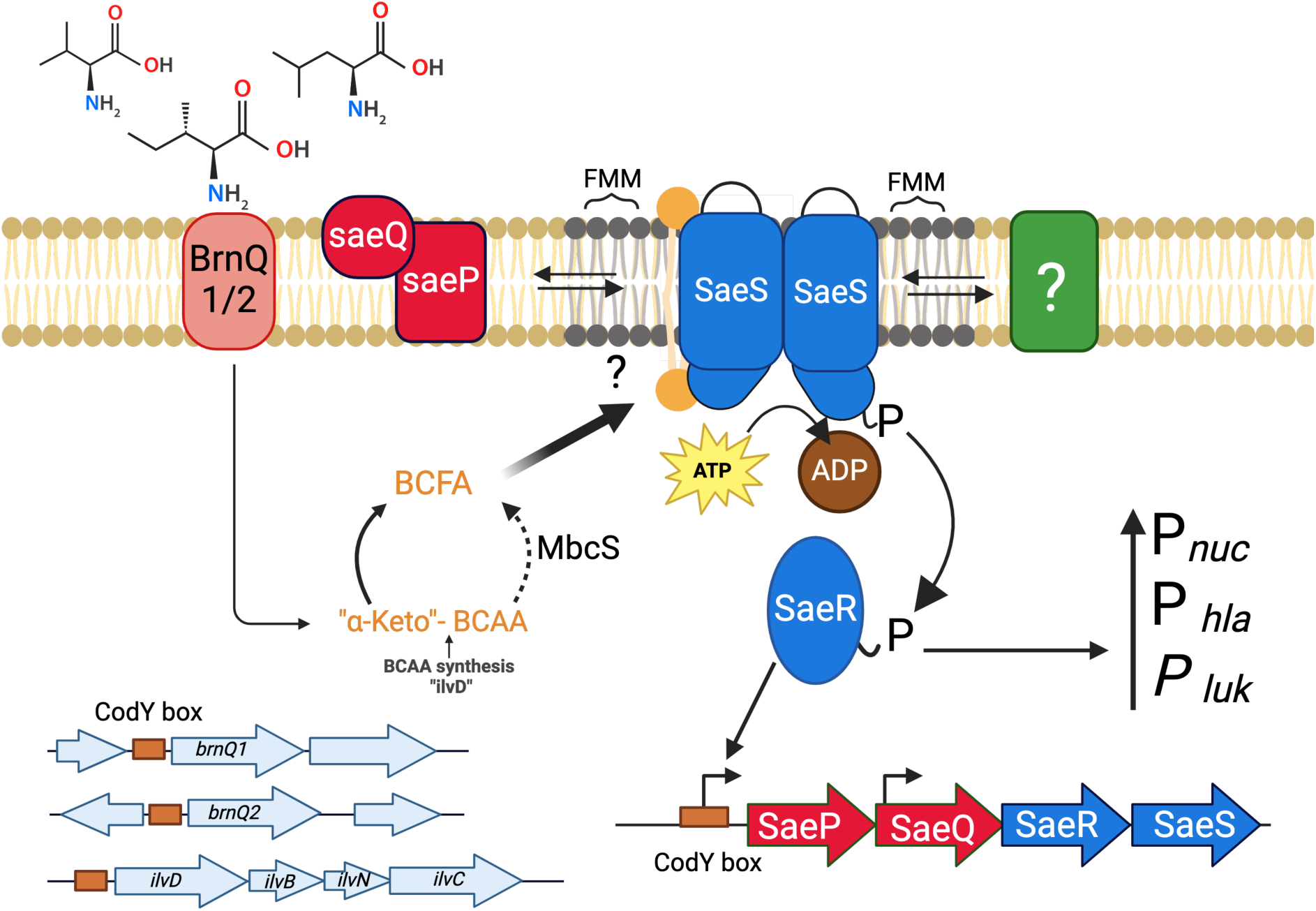
Working model for proposed mechanisms of SaeS activation by BCFA in *Staphylococcus aureus*. Under conditions of ILV limitation, the inactivation of CodY leads to the de-repression of genes *brnQ2* and the *ilv-leu* operon, enhancing the import and biosynthesis of isoleucine. Subsequent metabolic conversion of isoleucine results in the production of 15:0 and 17:0 anteiso BCFAs via BKDH-dependent and BKDH-independent pathways that upregulate SaeS kinase activity and increase cellular SaeR∼P levels. BCFA activation of SaeS may be direct or indirect altering protein-protein or protein-lipid interactions at the SaeS signaling complex. Green box, potential activator protein; solid line denotes BKDH-dependent BCFA synthesis pathway; dotted arrow denotes a second BKDH-independent and MbcS-dependent route to BCFA synthesis as described previously (55).

Lipids have also emerged as important regulatory effectors in prokaryotic and eukaryotic membrane receptors. Indeed, in addition to BCFAs, we have shown that cardiolipin is essential for full SaeS activity (60). BCFAs may bind directly to SaeS to facilitate dimerization, promote kinase activity, or act as an allosteric modulator to induce a conformational change. Alternatively, *a*15:0 may act indirectly either because a minor secondary metabolite of it interacts with SaeS or because it regulates a modification of SaeS. Nonnative mass spectrometry of SaeS would reveal the latter and the powerful new lipidomics techniques would reveal the former (61, 62). Thus, key lipids and membrane proteins may act as the true sensors for SaeS, and characterizing these interactions is a current focus of our lab. Because these new lipids and proteins are important for Sae TCS activity and virulence, they are potentially new anti-virulence targets. Answering these questions would provide insight into how *S. aureus* upregulates virulence factor production. Moreover, since other Gram-positive bacteria utilize IM-HKs, using SaeS as a model will provide insight into how these systems function generally.

## Materials and Methods

### Bacterial strains and growth conditions

All strains used in this study are derivatives of USA300 *S. aureus* LAC **(Table S1)**. *S. aureus* strains were cultured either in tryptic soy broth (TSB, Becton Dickenson formulation containing 2.5 g liter^-1^ dextrose) or chemically defined medium (CDM) (50). Briefly, CDM medium was formulated with alanine (672 μM), arginine (287 μM), aspartic acid (684 μM), cysteine (166 μM), glutamic acid (680 μM), glycine (670 μM), histidine (129 μM), isoleucine (228 μM), leucine (684 μM), lysine (342 μM), methionine (20 μM), phenylalanine (240 μM), proline (690 μM), serine (285 μM), threonine (260 μM), tryptophan (50 μM), tyrosine (275 μM), valine (684 μM), thiamine (56 μM), nicotinic acid (10 μM), biotin (0.04 μM), pantothenic acid (2.3 μM), MgCl_2_ (1,000 μM), CaCl_2_ (100 μM), monopotassium phosphate (40,000 μM), dipotassium phosphate (14,700 μM), sodium citrate dehydrate (1,400 μM), magnesium sulfate (400 μM), ammonium sulfate (7,600 μM), and glucose (27,753 μM). All strains were grown at 37°C. When necessary, media were solidified with agar (1.5% [wt / vol]) and supplemented with the following antibiotics at the indicated concentrations: chloramphenicol (Cm), 5 to 10 μg mL^-1^; erythromycin (Erm), 5 to 10 μg mL^-1^; or tetracycline (Tc) at 1.5 μg mL^-1^. When indicated, media were supplemented with 0.5 mM branched-chain carboxylic acids (*a*C_5_ [2-methylbutyric acid], *i*C_5_ [isovaleric acid], and *i*C_4_ [isobutyric acid]). Except for modifications pertinent to specific experiments described later, cultures utilized in experiments were grown overnight in test tubes from colonies picked from freshly streaked TSB-agar plates and then diluted 1:100 into flasks with the indicated medium (5:1 flask:medium volume). After 5-6 hours, cells were collected by centrifugation and used in experiments. Growth prior to the use of bacteria in individual experiments was monitored by the increase in absorbance at 600 nm (OD_600_) using an Amersham Ultraspec 2100 Pro UV-visible spectrophotometer.

### Genetic techniques

Oligonucleotides used in this study were synthesized by Integrated DNA Technologies (IDT; Coralville, IA) and are listed in **Table S2**. The Plasmids used in this study are listed in **Table S3**. Restriction enzymes, T4 DNA ligase, and Q5 DNA polymerase were purchased from New England Biolabs (NEB). Plasmid and genomic DNA (gDNA miniprep kits were purchased from Promega. *E. coli* NEB 5α (NEB) was used as a host for plasmids, which were then transferred when necessary into *S. aureus* strain RN4220 by electroporation as previously described (63). Plasmid and marked mutations were moved between *S. aureus* strains via Φ85- or Φ11-mediated transduction (64).

To construct the in-frame, Δ*ilvD* allele, ∼500 bp up- and down-stream of the *ilvD* coding sequence were amplified using *S. aureus* chromosomal DNA. A third, overlapping PCR was used to fuse the two DNA fragments. PCR reactions used oSRB368-371. This DNA fragment was cut and ligated to the same sites of pMAD, resulting in pSRB72. Allelic exchange was performed as described previously screening for white, Em^S^ colonies on TSB XG medium and ILV auxotrophy on CDM (65).

### Reporter fusion assays

Single colonies of *S. aureus* strains were freshly streaked out on plates from frozen stocks and were used to inoculate the appropriate medium. For experiments utilizing the Δ*lpdA mbcS1* Δ*codY* mutant strain, a mixture of the three branched-chain carboxylic acids (0.5 mM each) was added to the medium in each corresponding tube, and cultures were grown overnight before being diluted 1:5 in 100 μl total volume per well of 96-well plates. Both OD_600_ and GFP fluorescence (485 nm excitation, 535 nm emission) were read in a computer-controlled BioTek Synergy H1 Multimode plate reader (Agilent) using Gen5 IVD control software (version 1.11). Relative Fluorescence Units (RFU values) were calculated by dividing the 535 nm emission intensity values by the OD_600_ values. For experiments assaying the effect of branched-chain amino acid biosynthesis or import on Sae TCS activity, inoculated cultures were grown overnight and then diluted into fresh medium (with or without BCCAs, depending on the experiment) to an optical density (OD_600_) of 0.05, before 200 µl aliquots were transferred into each well of a 96-well plate. Both OD_600_ and GFP fluorescence were read every 20 minutes over 16 hours using the same BioTek Synergy H1 Multimode plate reader and RFU was calculated for each time point. Finally, for experiments involving overexpression of *brnQ2^+^*, the inoculated cultures were grown overnight in test tubes, then diluted 1:100 into flasks in media containing various concentrations of anhydrotetracycline. After 5-6 hours incubation at 37°C, 100 μl aliquots were transferred to wells of a 96-well plate and then read using the abovementioned plate reader to determine RFU.

### Nuclease activity quantification via FRET

Secreted nuclease activity was quantified using a fluorescence resonance energy transfer (FRET) assay described previously (66). In brief, cells were grown to an exponential phase in CDM. These cultures were then serially diluted and grown overnight. Cultures that reached an OD_600_ of ∼1 after overnight incubation were double back diluted to OD_600_ of 0.05 and allowed to grow back to the exponential phase (OD_600_ of 0.5). To obtain secreted nuclease, 0.7 mL of experimental culture was transferred to 0.22-μm Spin-X centrifuge tube filters (Corning Life Sciences) and centrifuged at 21,000 x g for 3 minutes. Sterilized culture supernatants were frozen at −20°C until use. The single-stranded oligonucleotide FRET substrate was diluted to a concentration of 2 mM in buffer A (20 mM Tris [pH 8.0], 0.5 M CaCl_2_) and then mixed 1:1 with thawed nuclease-containing supernatants that were diluted as described below. Fluorescence, indicative of the substrate cleavage by nuclease (535nm excitation, 590nm emission), was measured at 30°C using the BioTek Synergy H1 plate reader described above. The relative fluorescence units were converted to units of nuclease activity by interpolation using a standard curve generated with purified micrococcal nuclease enzyme (Worthington Biochemicals). To ensure that the readouts from the assay of the sterilized culture supernatants fell within the linear regression range of the standard curve samples, the nuclease-containing supernatants were diluted 1:10 in water and then serially diluted 2-fold in water up to 14 times before being mixed with FRET substrate.

### Membrane fatty acid analysis

Strains were grown in CDM medium to exponential phase, at which time a 15-ml sample of culture was pelleted (between 10-100 mg wet cell pellet) and washed twice with phosphate-buffered saline (PBS) before being stored at −80°C. The fatty acids in each pellet were saponified and methylated before being analyzed using gas chromatographic analysis of fatty acid methyl esters (GC-FAME) as a fee for service at the Center for Microbial Identification and Taxonomy (Norman, OK).

### Analysis of SaeS kinase activity

*In vivo* Separation of SaeR and SaeR∼P was performed as described previously using 12% polyacrylamide gels containing 100 µM Manganese and 50 µM of the acrylamide-pendant Phos-tag ligand (Pendleton 2022). Cells were grown in CDM to exponential phase (OD_600_ 0.5-1) in at 37°C with agitation. Cell pellets (OD_600_ of 10) were collected at 13,000 x g and stored at −80°C prior to analysis. Whole cell extracts were obtained by resuspending cell pellets in 300 µL cell extract buffer (20 mM Tris-HCl [pH 7.0], 1X Protease Inhibitor Cocktail Set I (Sigma-Aldrich)) and transferred to sterile screw cap tubes containing approximately 100 µL of 0.1mm silica beads. The cells were homogenized at room temperature using a Precellys 24 bead beater (Bertin technologies) for 3 cycles of 6500 rpm for 30 s each, followed by 3 mins pauses on ice. The tubes were then centrifuged at 8500 x g for 15 seconds to settle the beads, and the supernatant was transferred into new microcentrifuge tubes. Whole-cell extracts were normalized by protein concentration (A280) to 100 µg, mixed with 5X SDS loading dye, and electrophoresed on Phos-tag gels with standard running buffer (0.1% [wt/vol] SDS, 25 mM Tris-HCl, 192 mM glycine) at 4°C under constant voltage (150 V) for 2 h. The gels were washed for 15 minutes with transfer buffer (25 mM Tris [pH 8.3], 192 mM glycine, 20% methanol) with 1 mM EDTA followed by a second wash without EDTA to remove manganese ions. Proteins were then transferred to 0.45 µM PVDF membranes (Cytiva). Membranes were blocked in blocking buffer (5% [wt/vol] skim milk in Tris Buffered Saline with Tween 20 (TBST) (20 mM Tris-HCl, 150 mM NaCl with 0.1% [wt/vol] Tween 20 pH 7.6) for 1 h. Membranes were then subjected to three brief washes in TBST and incubated with polyclonal rabbit antibodies to SaeR (1.5:1,000) for 1 h. Membranes were then washed three times with and incubated with StarBright Blue 700 goat anti-rabbit IgG (1:3500; Bio-Rad) for 1 h. Membranes were subjected to three brief washes in TBST and signals were visualized using an Amersham ImageQuant800. The densities of the SaeR∼P relative to total SaeR signal were determined by quantification with Multi Gauge software (FujiFilm). The data are representative of three different independent experiments, and a representative image is shown.

### Analysis of SaeR and SrtA

The whole cell extracts and protein concentrations described above were used to visualize total SaeR levels. Briefly, 100µg of whole cell extracts mixed with 5X SDS loading dye were subjected to 12% SDS-PAGE, and proteins were transferred to 0.45 µM PVDF membranes (Cytiva). After transfer, membranes were blocked in blocking buffer for 1 h. Membranes were then washed three times with TBST and incubated with either polyclonal rabbit antibodies to SaeR (1.5:1000) or SrtA (1:1000) for 1 h. Membranes were then washed three times and incubated with StarBright Blue 700 goat anti-rabbit IgG (1:3500; Bio-Rad) for 1 h. Membranes were subjected to three brief washes in TBST, and signals were visualized using an Amersham ImageQuant800. The densities of the SaeR and SrtA signal were determined by quantification with Multi Gauge software (FujiFilm). The data are representative of three different independent experiments, and a representative image is shown.

## Acknowledgments

We thank Dr. David Heinrichs and Taeok Bae for the gift of plasmid pSO2 (*brnQ2^+^*) and SrtA and SaeR antibodies, respectively. We thank Deepak Sharma and Dr. François de Mets for technical assistance, and thank Dr. Paul Lawson and the Center for Microbial Identification and Taxonomy for GC-FAME analysis. We also thank members of the Georgetown University Microbial Interest Group as well as Brinsmade lab members for their helpful comments and discussions.

This work was supported in part by NIH grants R01 AI137403 and R00 GM099893 to SRB. The funders had no role in study design, data collection and interpretation, or the decision to submit the work for publication.

## Author Contributions

SRB conceived the study; SA and SRB conceptualized the research goals and aims; SA and DAD performed the investigations and analyzed the data, SRB, SA, and DAD prepared the original manuscript draft; SRB and SA prepared the final manuscript.

